# Rab12 regulates LRRK2 activity by promoting its localization to lysosomes

**DOI:** 10.1101/2023.02.21.529466

**Authors:** Vitaliy V. Bondar, Xiang Wang, Oliver B. Davis, Michael T. Maloney, Maayan Agam, Marcus Y. Chin, Audrey Cheuk-Nga Ho, David Joy, Joseph W. Lewcock, Gilbert Di Paolo, Robert G. Thorne, Zachary K. Sweeney, Anastasia G. Henry

**Affiliations:** Denali Therapeutics Inc., 161 Oyster Point Blvd., South San Francisco, CA 94080, USA; REGENXBIO Inc., 9804 Medical Center Drive, Rockville, MD 20850; Cellares, 341 Allerton Ave, South San Francisco, CA 94080; Department of Pharmaceutics, University of Minnesota, 308 Harvard Street SE, Minneapolis, MN 55455, USA; Interline Therapeutics Inc., 1400 Sierra Point Parkway, Suite 300 Brisbane, CA 94005, USA

## Abstract

Leucine-rich repeat kinase 2 (LRRK2) variants associated with Parkinson’s disease (PD) and Crohn’s disease lead to increased phosphorylation of its Rab substrates. While it has been recently shown that perturbations in cellular homeostasis including lysosomal damage and stress can increase LRRK2 activity and localization to lysosomes, the molecular mechanisms by which LRRK2 activity is regulated have remained poorly defined. We performed a targeted siRNA screen to identify regulators of LRRK2 activity and identified Rab12 as a novel modulator of LRRK2-dependent phosphorylation of one of its substrates, Rab10. Using a combination of imaging and immunopurification methods to isolate lysosomes, we demonstrated that Rab12 is actively recruited to damaged lysosomes and leads to a local and LRRK2-dependent increase in Rab10 phosphorylation. PD-linked variants, including LRRK2 R1441G and VPS35 D620N, lead to increased recruitment of LRRK2 to the lysosome and a local elevation in lysosomal levels of pT73 Rab10. Together, these data suggest a conserved mechanism by which Rab12, in response to damage or expression of PD-associated variants, promotes the recruitment of LRRK2 and phosphorylation of its Rab substrate(s) at the lysosome.

## Introduction

Coding variants in *LRRK2* can cause monogenic Parkinson’s disease (PD), and coding and noncoding variants in *LRRK2* are associated with increased risk for developing sporadic PD and Crohn’s disease (1–3). The majority of pathogenic LRRK2 variants cluster within its Roc-COR (Ras of complex proteins/C-terminal of Roc) GTPase tandem domain or kinase domain and contribute to disease risk by ultimately increasing LRRK2’s kinase activity (4–6). LRRK2 phosphorylates a subset of Rab GTPases, including Rab10 and Rab12. Excessive Rab phosphorylation can perturb interactions between Rabs and downstream effectors, which impairs various aspects of membrane trafficking including lysosomal function (7–9). LRRK2 localizes primarily to the cytosol in an inactive state, and higher-order oligomerization and membrane recruitment appear to be required for LRRK2 activation and Rab phosphorylation (10–13). Recent work suggests that interactions between LRRK2 and its phosphorylated Rab substrates help maintain LRRK2 on membranes and result in a cooperative, feed-forward mechanism to promote additional Rab phosphorylation (14). However, it is not clear what mechanisms promote the initial recruitment of LRRK2 to membranes to trigger Rab phosphorylation or whether increased LRRK2 membrane association is a common driver of the elevated LRRK2 activity observed in PD.

Endolysosomal genes that modify PD risk and lysosomal damage more generally observed in PD can also increase LRRK2 activation and phosphorylation of its Rab substrates(15–20). Rab29 and VPS35, proteins that are genetically-associated with PD and play key roles in lysosomal function by regulating sorting between the endolysosomal system and the trans-Golgi network, can modulate LRRK2 activity, as overexpression of Rab29 or expression of the pathogenic VPS35 D620N variant lead to significantly elevated LRRK2-mediated phosphorylation of Rab10 and other Rab substrates (18–20). LRRK2 kinase activity also appears to be increased in nonhereditary idiopathic PD patients (21–23), and emerging data suggest that lysosomal damage more broadly may be a common trigger for LRRK2 activation. Lysosomotropic agents that disrupt the endolysosomal pH gradient or puncture lysosomal membranes enhance LRRK2 recruitment to damaged lysosomes and result in increased Rab10 phosphorylation (15–17, 23). Several hypotheses around the purpose of LRRK2 recruitment to damaged lysosomes have been proposed, including promotion of lysosomal membrane repair or clearance of lysosomal content through exocytosis or sorting of vesicles away from damaged lysosomes (15–17). There is a clear need to better define how LRRK2 is recruited to lysosomes upon damage, how these steps translate to LRRK2 activation, and how broadly conserved these mechanisms are in PD.

Here, we identify Rab12 as a novel regulator of LRRK2 activity. Rab12 is required for LRRK2-dependent phosphorylation of Rab10 and mediates LRRK2 activation in response to lysosomal damage. We demonstrate that Rab12 promotes Rab phosphorylation by recruiting LRRK2 to lysosomes following lysosomal membrane rupture. Pathogenic PD variants including VPS35 D620N and LRRK2 R1441G also result in increased levels of LRRK2 and pT73 Rab10 on lysosomes. Together, our data delineate a conserved mechanism by which LRRK2 activity is regulated basally and in response to lysosomal damage and genetic variants associated with disease.

## Results

### siRNA-based screen identifies Rab12 as a key regulator of LRRK2 kinase activity

Although a subset of 14 Rab GTPases has been clearly established as LRRK2 substrates, increasing data suggest a reciprocal relationship exists in which Rab proteins may also contribute to LRRK2 membrane association and activation (14, 18, 20, 24). Previous studies have shown that overexpression of one such LRRK2 substrate, Rab29, can increase LRRK2-dependent phosphorylation of Rab10 by promoting its membrane association at the Golgi complex(18, 20, 24). However, additional work in RAB29 KO models demonstrated that LRRK2 activity was minimally impacted by loss of Rab29, suggesting Rab29 does not regulate LRRK2 activity under physiological conditions (25). To determine whether any LRRK2-Rab substrates were required for LRRK2 kinase activity, we performed a targeted siRNA screen on 14 Rab genes in human A549 cells that endogenously express both LRRK2 and Rab10. Rab10 phosphorylation was chosen as the endpoint to assess LRRK2 activation as it is an established readout of LRRK2 kinase activity that has been routinely used in preclinical and clinical settings (23, 26, 27). The levels of Rab10 phosphorylation were measured using a previously-described quantitative Meso Scale Discovery (MSD)-based assay (23). Greater than 50% knockdown of gene expression of each target was demonstrated for the targets assessed using qPCR-based analysis, and we confirmed that knockdown of the positive controls LRRK2 and RAB10 attenuated the phospho-Rab10 signal as expected (Figure 1A and B and Figure 1-figure supplement 1). RAB12 was the only hit gene whose knockdown significantly reduced Rab10 phosphorylation (Figure 1A). We confirmed that RAB12 knockdown reduced gene expression and led to a reduction in Rab12 protein levels using RAB12 KO A549 cells (Figure 1 C-E). RAB12 knockdown did not impact the levels of LRRK2 or Rab10, suggesting that Rab12 mediates Rab10 phosphorylation by regulating LRRK2’s activity rather than the stability of LRRK2 or Rab10 (Figure 1B,D and F and Figure 1-figure supplement 1).

**Figure 1:**
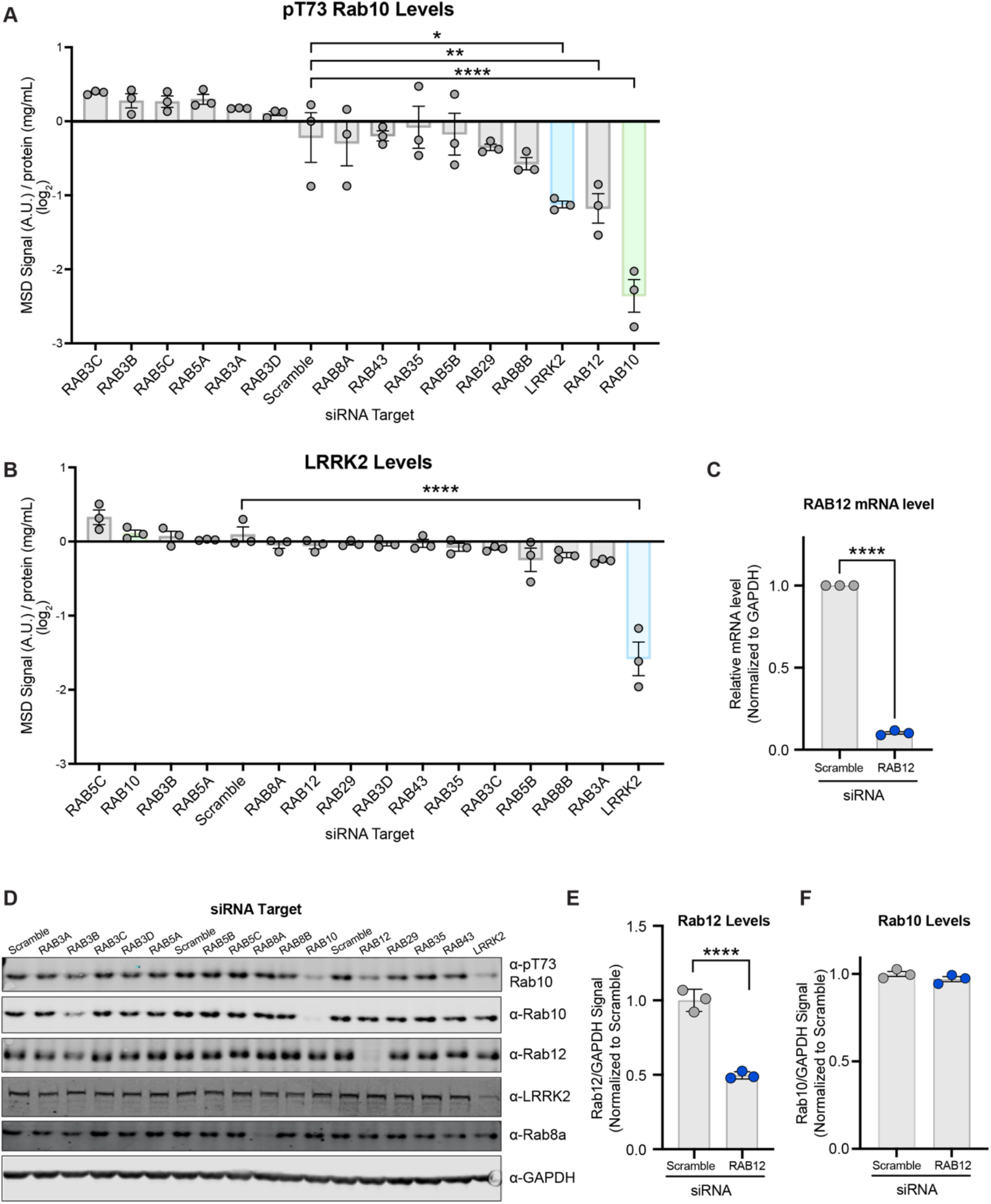
A targeted siRNA screen identifies Rab12 as a key regulator of LRRK2 kinase activity. A and B) A549 cells were transfected with siRNA targeting LRRK2 and its Rab substrates, lysed 3 days after transfection, and the levels of pT73 Rab10 and LRRK2 were quantified using MSD-based analysis. The MSD signal was normalized to the protein concentration, and data are shown on a log2 scale as the mean ± SEM; n=3 independent experiments, and statistical significance was determined using one-way ANOVA with Dunnett’s multiple comparison test. C) RAB12 mRNA levels were quantified using qPCR-based analysis and normalized to GAPDH following transfection with siRNAs targeting a scramble sequence or RAB12. Data are shown as the mean ± SEM; n=3 independent experiments, and statistical significance was determined using paired t-test. D) The levels of pT73 Rab10, Rab10, Rab12, LRRK2, and Rab8a following siRNA-mediated knockdown of LRRK2 and its Rab substrates were assessed in A549 cells by western blot analysis. Shown is a representative immunoblot with GAPDH as a loading control. E and F) Fluorescencesignals of immunoblots from multiple experiments were quantified, and the Rab12 and Rab10 signal was normalized to GAPDH, normalized to the median within each batch and expressed as a fold change compared to the scramble control; data are shown as the mean ± SEM; n=3 independent experiments. Statistical significance was determined using unpaired t-test. *p < 0.05, **p < 0.01, ****p < 0.0001

**Figure 1- figure supplement 1:**
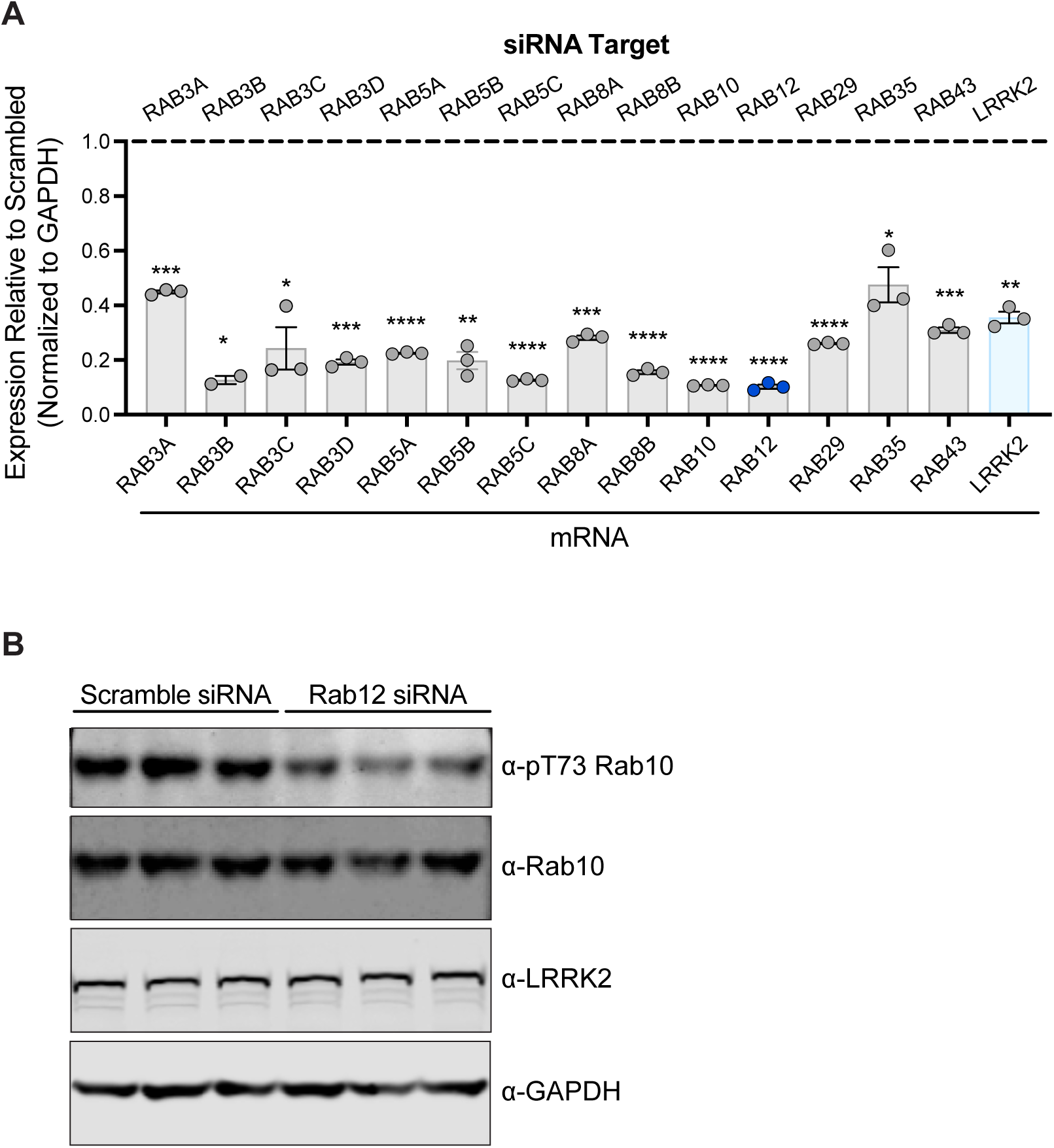
Confirmation of siRNA-mediated knockdown of LRRK2 Rab substrates. A) A549 cells were transfected with siRNA targeting LRRK2 and its Rab substrates, lysed 3 days after transfection, and knockdown was confirmed by qPCR-based analysis. The expression of each gene assessed was normalized to GAPDH expression, and then normalized to the expression observed with a scramble siRNA sequence. n=3 independent experiments. Data are shown as the mean ± SEM, with p values based on paired t-test. B) A549 cells were transfected with a scramble siRNA sequence or siRNA targeting RAB12, and the levels of pT73 Rab10, Rab10, and LRRK2 were assessed by western blot analysis. GAPDH was used as a loading control. *p < 0.05, **p < 0.01, ***p < 0.001****p < 0.0001

### Rab12 is required for LRRK2-dependent phosphorylation of Rab10

To confirm that Rab12 regulates LRRK2-dependent Rab10 phosphorylation, we used RAB12 KO A549 cells to assess the impact of RAB12 deletion using MSD- and western-blot analysis. We demonstrated that loss of Rab12 significantly impairs Rab10 phosphorylation at T73, showing a comparable reduction to that observed with loss of LRRK2 (Figure 2A and B and Figure 2- figure supplement 1). Total Rab10 levels were not reduced with RAB12 deletion and, in fact, were elevated in two out of three RAB12 KO clones assessed, confirming that the impact of loss of Rab12 on Rab10 phosphorylation cannot be explained by an effect on the protein levels of Rab10 (Figure 2C). Rab12 is itself a substrate for LRRK2, and we next explored whether LRRK2-mediated phosphorylation of Rab12 contributed to LRRK2 activation. To assess this, we generated doxycycline-inducible stable cell lines in the RAB12 KO cell background to allow overexpression of wildtype (WT) or a phospho-deficient mutant Rab12 in which the LRRK2 phosphorylation site (S106) was converted to an alanine (Figure 2D). Overexpression of either WT Rab12 or Rab12 S106A restored Rab10 phosphorylation at T73 and did not impact LRRK2 levels (Figure 2E and F). This finding was further confirmed using Rab12 S106A KI cells generated using CRISPR-Cas9 (Figure 2- figure supplement 1).

**Figure 2:**
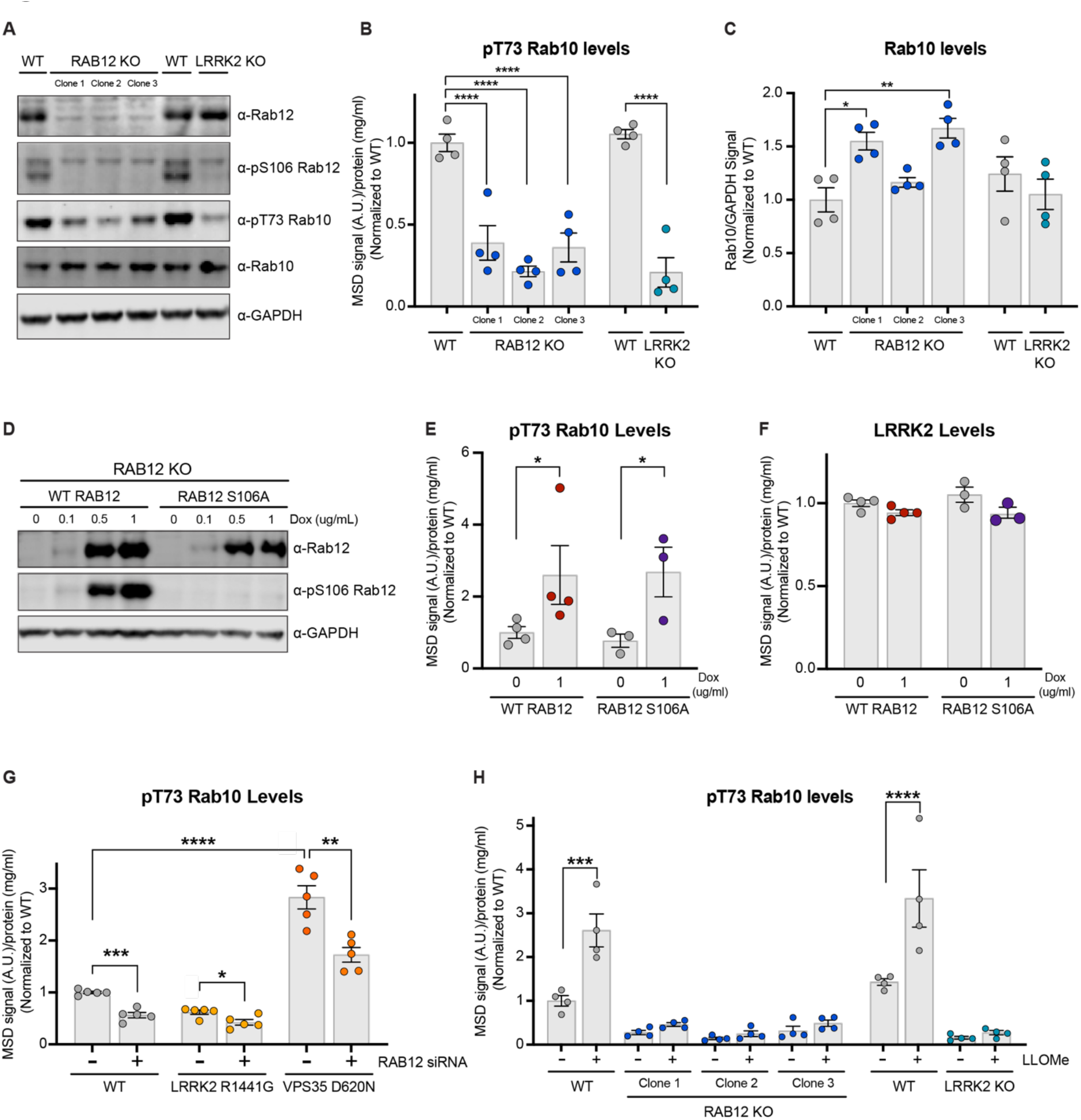
Rab12 is required for LRRK2 activation basally and in response to lysosomal damage and genetic variants associated with PD. **A)** The levels of Rab12, pS106 Rab12, pT73 Rab10, and Rab10 were assessed in WT, RAB12 KO, and LRRK2 KO A549 cells by western blot analysis. Shown is a representative immunoblot with GAPDH as a loading control. B) The levels of pT73 Rab10 were measured using an MSD-based assay. The MSD signal was normalized for protein input and expressed as a fold change compared to WT A549 cells; data are shown as the mean ± SEM; n=4 independent experiments, and statistical significance was determined using one-way ANOVA with Dunnett’s multiple comparison test. C) Fluorescence signals of immunoblots from multiple experiments were quantified, and the Rab10 signal was normalized to GAPDH and expressed as a fold change compared to WT A549 cells. Data are shown as the mean ± SEM; n=4 independent experiments. Statistical significance was determined using one-way ANOVA with Dunnett’s multiple comparison test. D-F) RAB12 KO A549 cells with doxycycline-inducible expression of WT RAB12 or a phospho-deficient variant of RAB12 (S106A) were treated with increasing concentrations of doxycycline for three days, and the levels of Rab12, pS106 Rab12, pT73 Rab10, and LRRK2 were measured. D) A representative immunoblot is shown assessing Rab12 and pS106 Rab12 protein levels following doxycycline-induced expression of WT or RAB12 S106A, and GAPDH was used as a loading control. E and F) The levels of pT73 Rab10 and LRRK2 were measured using MSD-based assays. MSD signals were normalized for protein concentration, and data were then normalized to the median within each batch and to the signals from the control group (RAB12 KO cells with inducible expression of WT Rab12 without doxycycline treatment). Data are shown as mean ± SEM; n=3-4 independent experiments, and statistical significance was determined using unpaired t-test on log transformed data. G) The impact of Rab12 knockdown was measured in WT, LRRK2 R1441G KI, and VPS35 D620N KI A549 cells. Cells were transfected with siRNA targeting RAB12, and pT73 Rab10 levels were measured by MSD-based analysis three days after transfection. The MSD signal was normalized for protein input and then normalized to the median within each batch and is expressed as a fold change compared to WT A549 cells transfected with scramble siRNA. Data are shown as the mean ± SEM; n=5 independent experiments. Statistical significance was determined using oneway ANOVA with Tukey’s multiple comparison test on log transformed data. H) WT, RAB12 KO, and LRRK2 KO A549 cells were treated with vehicle or LLOMe (1mM) for 2 hours, and the impact of LLOMe treatment on pT73 Rab10 levels was measured by MSD-based analysis. The MSD signal was normalized for protein input and is expressed as a fold change compared to WT A549 cells treated with vehicle. Data are shown as the mean ± SEM; n=4 independent experiments. Statistical significance was determined using two-way ANOVA with Sidak’s multiple comparison test. *p < 0.05, **p < 0.01, ***p < 0.001****p < 0.0001

**Figure 2- figure supplement 1:**
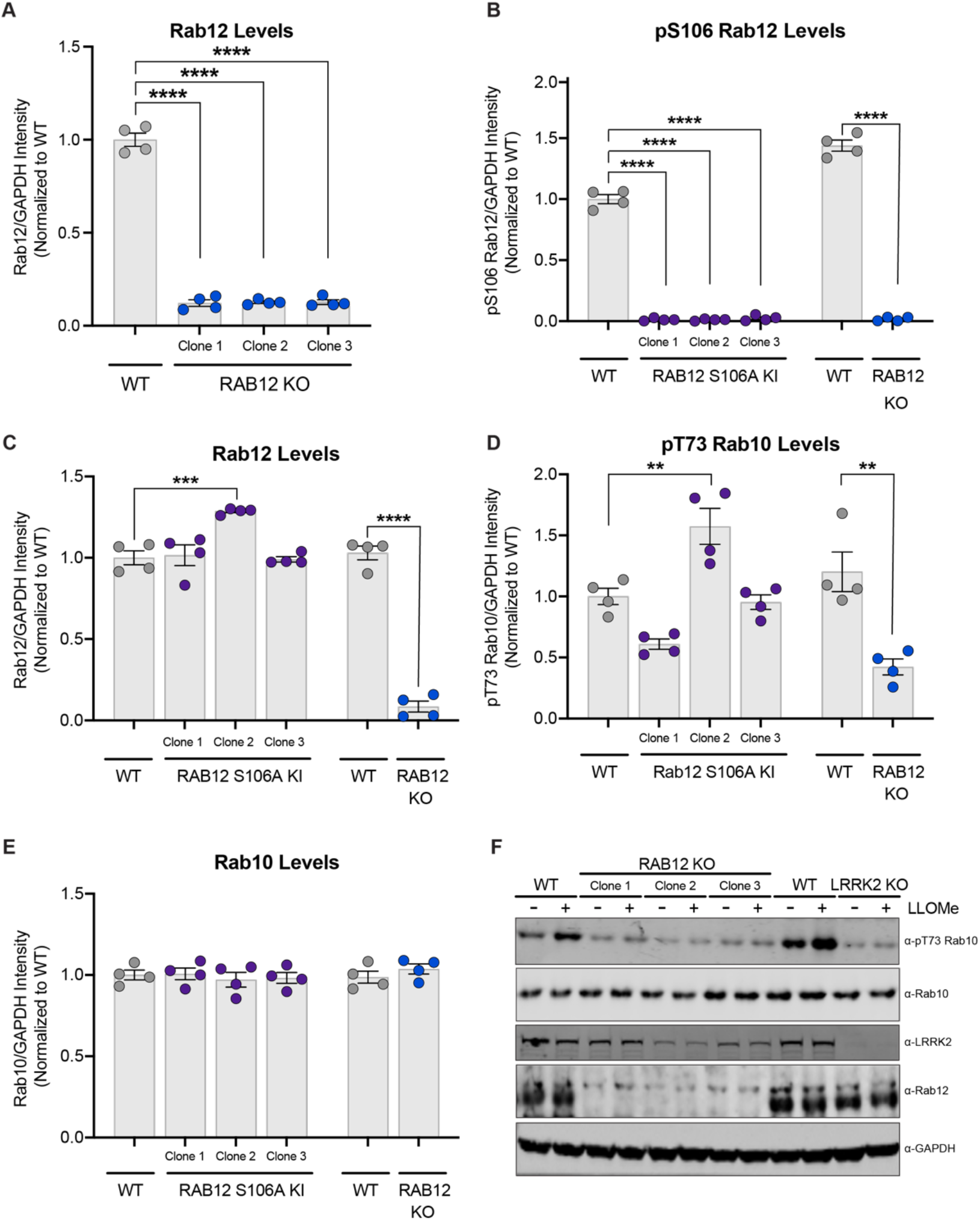
Confirmation of effects of RAB12 KO and RAB12 S106A KI on total and phospho-Rab12 and Rab10. A) The levels of Rab12 were measured in cell lysate from WT or three clones of RAB12 KO A549 cells by western blot analysis. The Rab12 signal was quantified and normalized to the GAPDH signal and expressed as a fold change compared to WT cells. Data are shown as the mean ± SEM; n=4 independent experiments. Statistical significance was determined using oneway ANOVA with Dunnett’s multiple comparisons test. B-E) The levels of pS106 Rab12, Rab12, pT73 Rab10, and Rab10 were measured in cell lysates from WT, three clones of RAB12 S106A KI A549, or RAB12 KO cells by western blot analysis, and GAPDH was used as a loading control. The pS106 Rab12 signal (B), Rab12 signal (C), pT73 Rab10 signal (D) and Rab10 signal (E) were measured and normalized to the GAPDH signal and expressed as a fold change compared to WT cells. Data are shown as the mean ± SEM; n=4 independent experiments. Statistical significance was determined using one-way ANOVA with Dunnett’s multiple comparison test. F) WT, RAB12 KO, and LRRK2 KO A549 cells were treated with vehicle or LLOMe (1 mM) for 2 hours, and the impact of LLOMe treatment on the levels of pT73 Rab10, Rab10, LRRK2, and Rab12 was assessed by western blot analysis. GAPDH was used as a loading control. **p < 0.01, ***p < 0.001****p < 0.0001

### Rab12 promotes LRRK2 activation by PD-linked genetic variants or lysosomal damage

Previous studies have established that pathogenic PD-linked variants and lysosomal membrane disruption can lead to increased LRRK2 kinase activity (15, 16, 19, 23). We next explored whether Rab12 might mediate LRRK2 activity in the context of either a pathogenic variant in *LRRK2* (LRRK2 R1441G) or *VPS35* (D620N KI). Unlike LRRK2 variants that lie within its kinase domain and directly increase kinase activity, including the common G2019S variant, the R1441G variant resides within the GTPase domain of LRRK2. The mechanisms by which the LRRK2 R1441G impact its kinase activity remain poorly defined. Rab10 phosphorylation was significantly reduced with RAB12 knockdown in LRRK2 R1441G KI and VPS35 D620N KI A549 cells (Figure 2G). Lysosomal membrane damage also increases LRRK2’s kinase activity, as treatment with L-leucyl-L-Leucine methyl ester (LLOMe), a lysosomotropic agent that condenses into membranolytic polymers and ruptures the lysosomal membrane, has been shown to increase LRRK2-dependent phosphorylation of its Rab substrates (15, 16). We confirmed that LLOMe treatment led to a significant increase in Rab10 phosphorylation in WT cells and demonstrated this effect was abolished in RAB12 KO cells (Figure 2H and Figure 2- figure supplement 1). Together, these data demonstrate that Rab12 is required to mediate LRRK2 activation in response to specific genetic variants associated with PD and lysosomal stress more broadly.

### Rab12 mediates LRRK2-dependent phosphorylation of Rab10 on lysosomes

Lysosomal membrane permeabilization has been shown to increase the levels of LRRK2 and pT73 Rab10 associated with lysosomes using overexpression systems (15, 16). Our data suggested that Rab12 may play a key role in promoting the recruitment of LRRK2 and ultimate phosphorylation of Rab10 on lysosomes in response to lysosomal damage. To assess this, we employed an established lysosome immunoprecipitation (Lyso-IP) method that enables the rapid isolation of lysosomes to determine whether Rab12 was required for Rab10 phosphorylation on lysosomes under endogenous expression conditions (28). Lysosomes isolated from WT and RAB12 KO A549 cells treated with vehicle or LLOME (1mM) for 2 hours displayed increased levels of endogenously-expressed galectin-3 (Gal3), confirming LLOMe treatment induced lysosomal membrane rupture and exposed luminal beta-galactosides (Figure 3A)(29–31).

**Figure 3:**
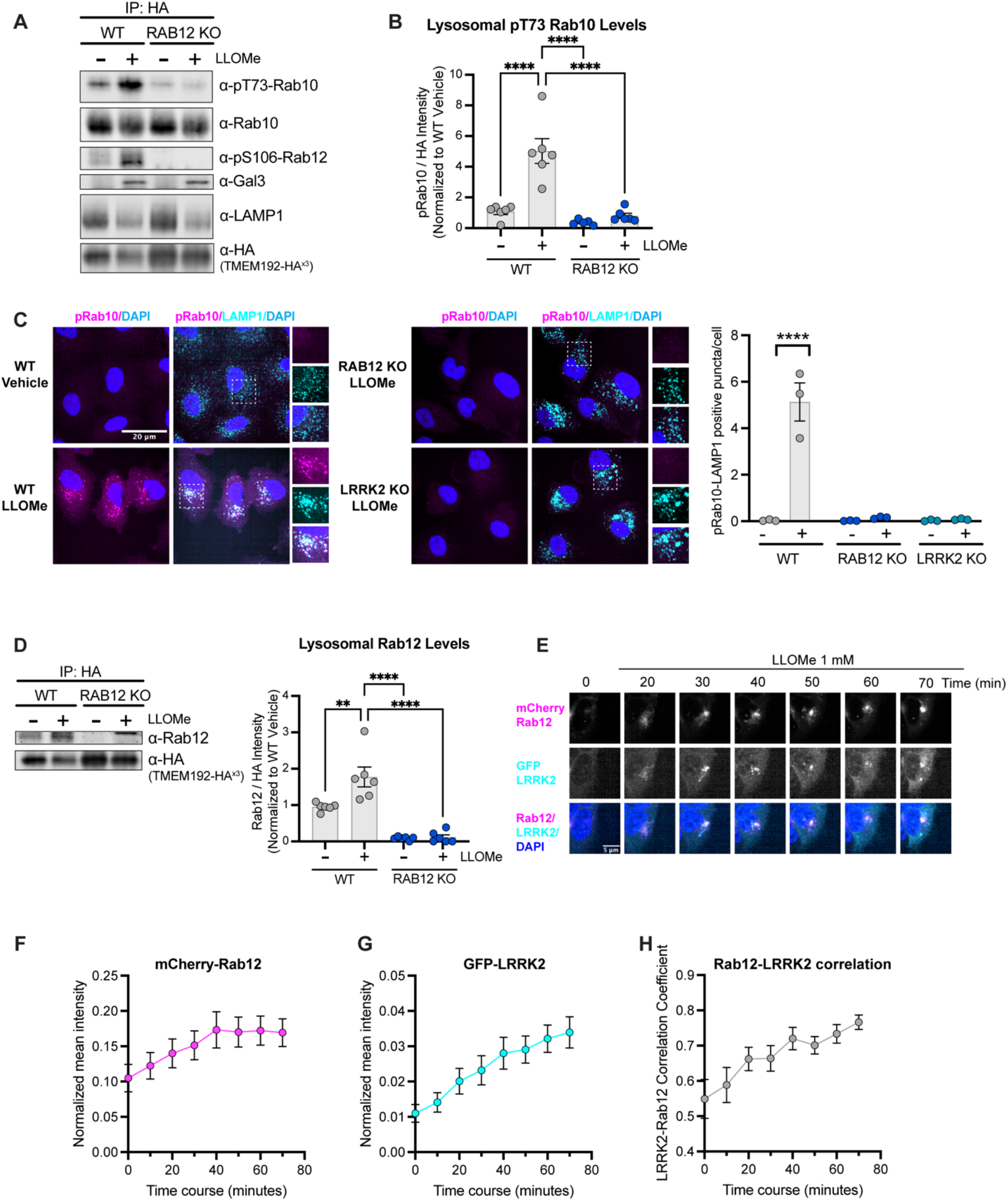
Rab12 is recruited to lysosomes following lysosomal damage and promotes Rab10 phosphorylation at the lysosome. A-B) Lysosomes were isolated from WT and RAB12 KO A549 cells treated with vehicle or LLOMe (1 mM) for 2 hours. The levels of pT73 Rab10, total Rab10, pS106 Rab12, Gal3, LAMP1 and HA were assessed by western blot analysis, and shown is a representative immunoblot. Fluorescence signals of immunoblots from multiple experiments were quantified, normalized to the median within each experimental replicate, and the pT73 Rab10 signal was normalized to the HA signal and expressed as a fold changed compared to lysosomes isolated from WT A549 cells treated with vehicle. n=6 independent experiments. Data are shown as the mean ± SEM, and statistical significance was determined using one-way ANOVA with Tukey’s multiple comparison test. C) WT, RAB12 KO and LRRK2 KO A549 cells were treated with vehicle or LLOMe (1 mM) for 2 hours, and the signals of pT73 Rab10 and LAMP1 were assessed by immunostaining. Scale bar, 20 μm. pT73 Rab10 (shown in magenta) and LAMP1 (shown in cyan) double positive puncta (i.e. overlap of magenta and cyan and shown in white) were quantified per cell from n = 3 independent experiments. Data are shown as the mean ± SEM with and statistical significance was determined using two-way ANOVA with Sidak’s multiple comparison test. D) Lysosomal Rab12 levels were assessed by western blot analysis from lysosomes isolated from WT and RAB12 KO A549 cells treated with vehicle or LLOMe (1 mM) for 2 hours. The Rab12 signals were normalized to the median within each experimental replicate, and then to the HA signal and expressed as a fold changed compared to lysosomes isolated from WT A549 cells treated with vehicle. n=6 independent experiments. Data are shown as the mean ± SEM with and statistical significance was determined using one-way ANOVA with Tukey’s multiple comparison test. E) Representative live-cell images show the recruitment of Rab12 and LRRK2 upon LLOMe (1 mM) treatment. HEK293 cells were transfected with mCherry-Rab12 and eGFP-LRRK2, and images were acquired every 10 min. Scale bar, 5 μm. F-G) Normalized mean intensity of mCherry-Rab12 (F) and eGFP-LRRK2 (G) were quantified over time in segmented cells (n=24 cells). H) Pearson’s correction coefficient (PCC) between normalized GFP-LRRK2 and mCherry-Rab12 were quantified over time in segmented cells (n=24 cells). n=3 experiments. Data are shown as mean ± SEM. **p < 0.01 and ****p < 0.0001

LLOMe treatment increased phosphorylation of Rab10 on isolated lysosomes from WT cells but failed to increase Rab10 phosphorylation on lysosomes from RAB12 KO cells, demonstrating that Rab12 is a critical regulator of Rab10 phosphorylation on lysosomes following lysosomal damage (Figure 3A and B and Figure 3- figure supplement 1). To explore this effect further, we visualized phosphorylated Rab10 on lysosomes following LLOMe treatment in WT and RAB12 KO cells. LLOMe treatment significantly increased colocalization between pT73 Rab10 and the lysosomal marker lysosomal-associated membrane protein 1 (LAMP1) in WT cells but had no effect in RAB12 KO cells, confirming that Rab12 is required for Rab10 phosphorylation on lysosomes in response to membrane rupture (Figure 3C and Figure 3- figure supplement 1).

**Figure 3- figure supplement 1:**
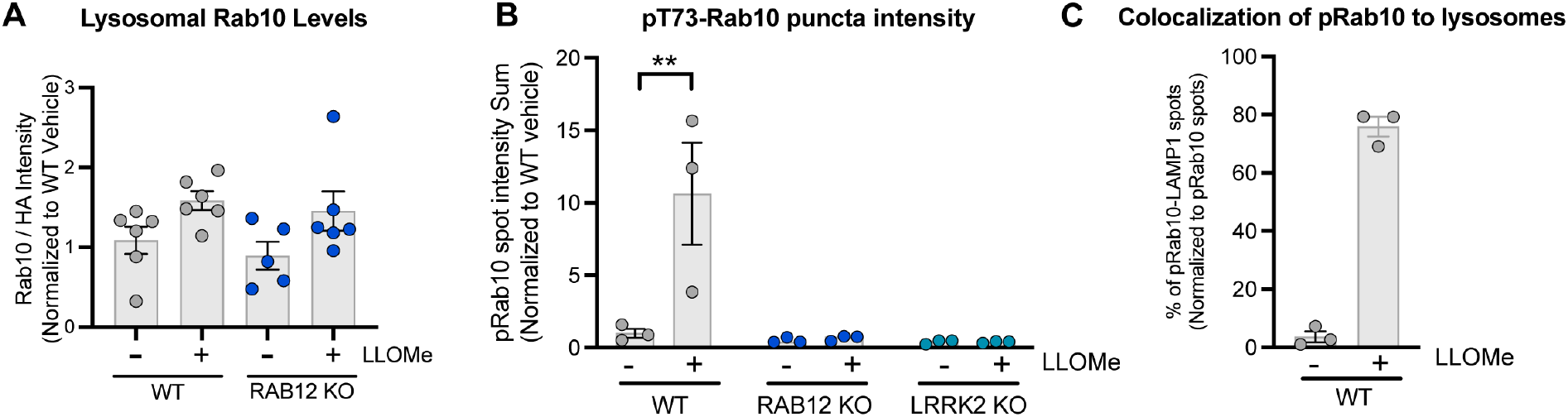
Analysis of Rab10 levels on the lysosomes and colocalization analysis of pT73 Rab10 with lysosomes in response to lysosomal damage. **A)** Lysosomal Rab10 levels were assessed by western blot analysis from lysosomes isolated from WT and RAB12 KO A549 cells treated with vehicle or LLOMe (1 mM) for 2 hours. All band intensities were normalized to the median within each experimental replicate, and Rab10 levels were then normalized to the HA signals and expressed as a fold change compared to lysosomes isolated from WT cells treated with vehicle (corresponding to Figure 3A); n=6 independent experiments. B) pT73 Rab10 signals were assessed by immunostaining of WT, RAB12 KO and LRRK2 KO A549 cells treated with vehicle or LLOMe (1 mM) for 2 hours (corresponding to Figure 3C). The sum intensity of puncta per cell was quantified from n=3 independent experiments. Data are shown as the mean ± SEM, and statistical significance was determined using two-way ANOVA with Sidak’s multiple comparison test. C) Percentage of pT73 Rab10 puncta colocalized with LAMP1 were quantified from WT A549 cells treated with vehicle or LLOMe (1 mM) for 2 hours. Data are shown as the mean ± SEM, n=3 independent experiments. **p < 0.01

### Rab12 increases Rab10 phosphorylation by promoting LRRK2 recruitment to lysosomes

We hypothesized that lysosomal membrane permeabilization may increase Rab12 recruitment to damaged lysosomes and that increased Rab12 levels on lysosomes may mediate the enhanced association of LRRK2 with lysosomes upon damage. Consistent with this idea, treatment with LLOMe significantly increased the levels of Rab12 on isolated lysosomes assessed by western blot analysis (Figure 3D). To gain additional insight around the dynamics of Rab12 and LRRK2 recruitment following lysosomal membrane permeabilization, we performed live-cell imaging of HEK293 cells overexpressing mCherry-tagged Rab12 and eGFP-tagged LRRK2 and assessed Rab12 and LRRK2 localization over time. Rab12 and LRRK2 showed a primarily cytosolic localization under baseline conditions, while LLOMe treatment increased the recruitment of Rab12 and LRRK2 to vesicular structures. The vesicular recruitment of Rab12 was observed as early as 20 minutes following LLOMe treatment and showed maximal recruitment at 40 minutes after LLOMe addition (Figure 3E-G). LRRK2 redistributed more slowly following LLOMe treatment, with its vesicular localization continuing to increase over the course of the experiment (Figure 3E and G). Rab12 colocalization with LRRK2 increased over time following LLOMe treatment, supporting a potential interaction between Rab12 and LRRK2 on lysosomes upon damage (Figure 3H). Together, these data demonstrate that Rab12 and LRRK2 both associate with lysosomes following membrane rupture and suggest that Rab12 recruitment may precede that of LRRK2.

To more directly assess whether Rab12 mediates the recruitment of LRRK2 to lysosomes upon lysosomal damage, we next examined the impact of RAB12 deletion on the lysosomal recruitment of LRRK2 upon lysosomal membrane permeabilization. Lysosomes were isolated from WT and RAB12 KO cells, and the endogenous levels of LRRK2 on lysosomes were quantified by western blot analysis. Lysosomal levels of LRRK2 were increased by approximately 2.5-fold following treatment with LLOMe in WT cells, and loss of Rab12 abrogated this LLOMe-induced increase (Figure 4A). These data show that Rab12 is required to promote LRRK2 localization to lysosomes following membrane damage.

**Figure 4:**
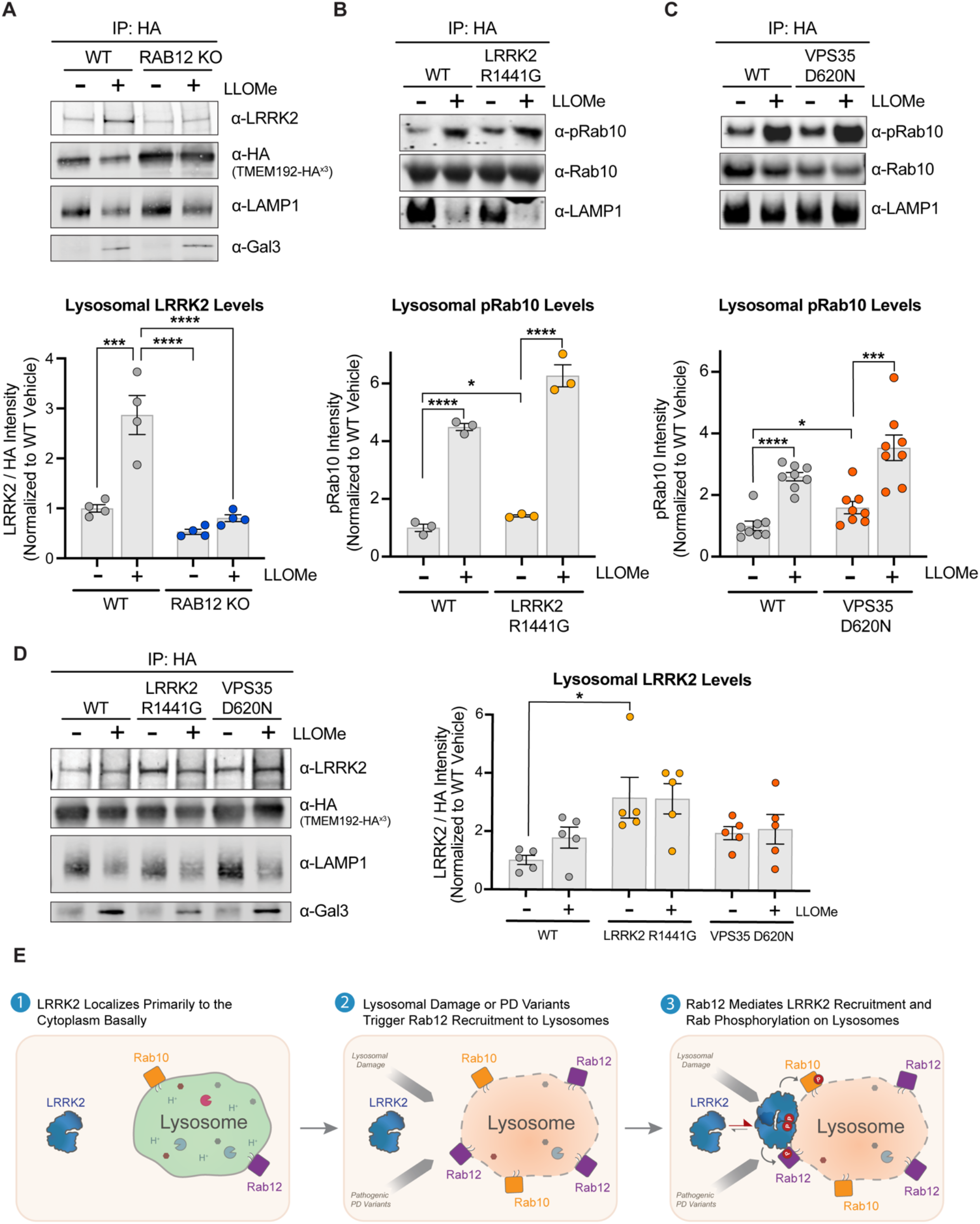
LRRK2 levels are increased on lysosomes following lysosomal damage in a Rab12-dependent manner and are also increased by PD-linked variants. A) To analyze lysosomal LRRK2 levels, lysosomes were isolated from WT and RAB12 KO A549 cells treated with vehicle or LLOMe (1mM) for 4 hours. The levels of LRRK2, HA, LAMP1 and Gal3 were assessed by western blot analysis, and shown is a representative immunoblot. Fluorescence signals of immunoblots from multiple experiments were quantified, normalized to the median within each experiment, and the LRRK2 signal was normalized to the HA signal and expressed as a fold change compared to lysosomes isolated from WT A549 cells treated with vehicle. Data are shown as the mean ± SEM; n=4 independent experiments. Statistical significance was determined using one-way ANOVA with Tukey’s multiple comparison test with a single pooled variance. B-C) To analyze lysosomal pRab10 levels, lysosomes were isolated from WT and LRRK2 R1441G KI (B) or VPS35 D620N KI (C) A549 cells treated with vehicle or LLOMe (1mM) for 2 hours, and the levels of pT73 Rab10, Rab10, and LAMP1 were assessed by western blot analysis. Fluorescence signals of immunoblots from multiple experiments were quantified, and the pT73 Rab10 signal was expressed as a fold change compared to lysosomes isolated from WT A549 cells treated with vehicle. Data are shown as the mean ± SEM; n=3 independent experiments (B) and n=8 independent experiments (C). Statistical significance was determined using unpaired t-test. D) Lysosomes were isolated from WT, LRRK2 R1441G KI, and VPS35 D620N KI A549 cells treated with vehicle or LLOMe (1mM) for 4 hours. The levels of LRRK2, HA, LAMP1, and Gal3 were assessed by western blot analysis and shown is a representative immunoblot. Fluorescence signals of immunoblots from multiple experiments were quantified, normalized to the median within each experiment, and the LRRK2 signal was normalized to the HA signal and expressed as a fold change compared to lysosomes isolated from WT A549 cells treated with vehicle. Data are shown as the mean ± SEM; n=5 independent experiments. Statistical significance was determined using one-way ANOVA with Dunnett’s multiple comparison test. *p < 0.05, ***p < 0.001****p < 0.0001 E) Model for proposed mechanism by which Rab12 promotes LRRK2 activation. Under steadystate conditions, LRRK2 localizes primarily to the cytoplasm. Lysosomal damage prompts the recruitment of Rab12, and Rab12 mediates the recruitment of LRRK2 to damaged lysosomes. An elevated local concentration of LRRK2 on lysosomes increases the likelihood for interactions with Rab GTPases localized on the lysosomal membrane, promoting LRRK2-dependent phosphorylation of its Rab substrates.

Enhanced recruitment of LRRK2 to lysosomes may promote Rab phosphorylation by effectively increasing the local concentration of LRRK2 in proximity to its Rab substrates, and we hypothesized that such a mechanism might explain LRRK2 activation observed in additional contexts beyond lysosomal damage. We next examined whether two PD-linked genetic variants associated with increased LRRK2 kinase activity, LRRK2 R1441G and VPS35 D620N, also had increased levels of LRRK2 on lysosomes. Lysosomes were isolated from WT, LRRK2 R1441G KI, and VPS35 D620N KI A549 cells at baseline and following LLOMe treatment, and the levels of total and phospho-Rab10 and LRRK2 were subsequently assessed by western blot analysis. Expression of LRRK2 R1441G or VPS35 D620N led to an increase in Rab10 phosphorylation on isolated lysosomes at baseline (Figure 4 B and C). The levels of LRRK2 on lysosomes were significantly increased in untreated LRRK2 R1441G KI cells and showed a trend for elevation in the VPS35 D620N KI cells (Figure 4D). In contrast to the effect observed in WT cells, LLOMe treatment failed to further increase LRRK2 levels in LRRK2 R1441G and VPS35 D620N cells. This result suggests that these pathogenic variants may have already maximally engaged this mechanism and are unable to further respond to additional lysosomal stress.

## Discussion

Increased LRRK2 kinase activity is observed with pathogenic PD-linked variants and in sporadic PD patients, but many questions remain regarding the mechanisms by which LRRK2 activation is regulated basally and in response to endolysosomal stress associated with disease. We identify Rab12 as a novel regulator of LRRK2 activation and demonstrate that Rab12 plays a critical role in mediating LRRK2-dependent Rab phosphorylation in response to lysosomal damage. Our data show that lysosomal membrane rupture promotes Rab12 localization to lysosomes. We propose a model in which Rab12 recruits LRRK2 to the lysosome and enhances its activity on lysosomal membranes by increasing LRRK2’s local concentration near Rab10 and potentially other Rab substrates (Figure 4E). Previous studies suggested that another LRRK2 substrate, Rab29, can regulate LRRK2’s kinase activity and showed that exogenously-expressed Rab29 is capable of activating LRRK2 at the trans-Golgi or at additional membranes when artificially-anchored there (20, 24). However, these studies relied on significant overexpression of Rab29 to increase LRRK2 activity, and analyses of Rab29 KO cellular models or mice showed that RAB29 deletion minimally impacted LRRK2-dependent Rab10 phosphorylation (25). Here, we used endogenous expression conditions to demonstrate that Rab12, but not Rab29, is required for LRRK2-mediated phosphorylation of Rab10 and that Rab12 mediates the localization and activation of LRRK2 on lysosomes upon lysosomal stress. The PD-linked pathogenic LRRK2 variant R1441G, which lies outside of LRRK2’s kinase domain, and VPS35 D620N variant have been shown in previous studies and confirmed here to increase LRRK2 activity (18–20). Our results suggest that these variants may also promote LRRK2 activation by increasing Rab12-mediated LRRK2 localization to lysosomes, implying this may represent a general mechanism by which LRRK2 activity is increased by various genetic variants and environmental stressors associated with PD. Additional studies are warranted to determine how lysosomal membrane rupture triggers Rab12 recruitment and to better define how broadly such mechanisms are employed to drive LRRK2 activation in PD.

Our findings provide key insight into the mechanism by which LRRK2 activity is increased in response to lysosomal damage by demonstrating that Rab12 mediates LRRK2 localization to ruptured lysosomes. While the purpose of LRRK2 recruitment and activation on lysosomes is poorly defined, several hypotheses have been proposed suggesting that LRRK2 activity may be upregulated as a compensatory response to restore lysosomal homeostasis following damage. Previous work supports a model in which LRRK2 activity controls the decision to repair minor damage to lysosomal membranes through an ESCRT-mediated process or to target damaged lysosomes for destruction via lysophagy (17). Additional studies suggested LRRK2 hyperactivation triggers mechanisms aimed at clearing cargo from damaged lysosomes that have lost their proteolytic capacity, either through direct exocytosis of lysosomes or through a novel sorting process termed LYTL (lysosomal tubulation/sorting driven by LRRK2) in which tubules bud off from lysosomes following membrane rupture (15, 16, 32). LRRK2 activation in the face of minor or acute lysosomal insults likely plays a beneficial role to maintain lysosomal function, while chronic LRRK2 activation triggered by genetic variants with increased kinase activity or low-level lysosomal damage over time may ultimately impair the ability to effectively respond to membrane stress and maintain the balance between lysosomal repair and destruction. Macrophages from PD patients carrying pathogenic LRRK2 variants were shown to accumulate more damaged lysosomes compared to samples from healthy controls, suggesting that mechanisms that respond to and repair lysosomal membrane rupture may be perturbed in PD(17). A deeper understanding of the mechanisms by which LRRK2 responds to lysosomal damage and how these contribute to PD pathogenesis is critical to guide new potential therapeutic strategies targeting LRRK2 for the treatment of PD.

## Materials and Methods

### Generation of CRISPR KO and KI cell lines

Cell line engineering of A549 cells to generate homozygous LRRK2 R1441G (CGC/GGC) knockin, homozygous LRRK2 KO, homozygous RAB12 KO, and homozygous VPS35 D620N (GAT/ATT) knock-in was performed using CRISPR/Cas9. Sequence information for generating targeting gRNA, ssODN donor and PCR primers are as follows:

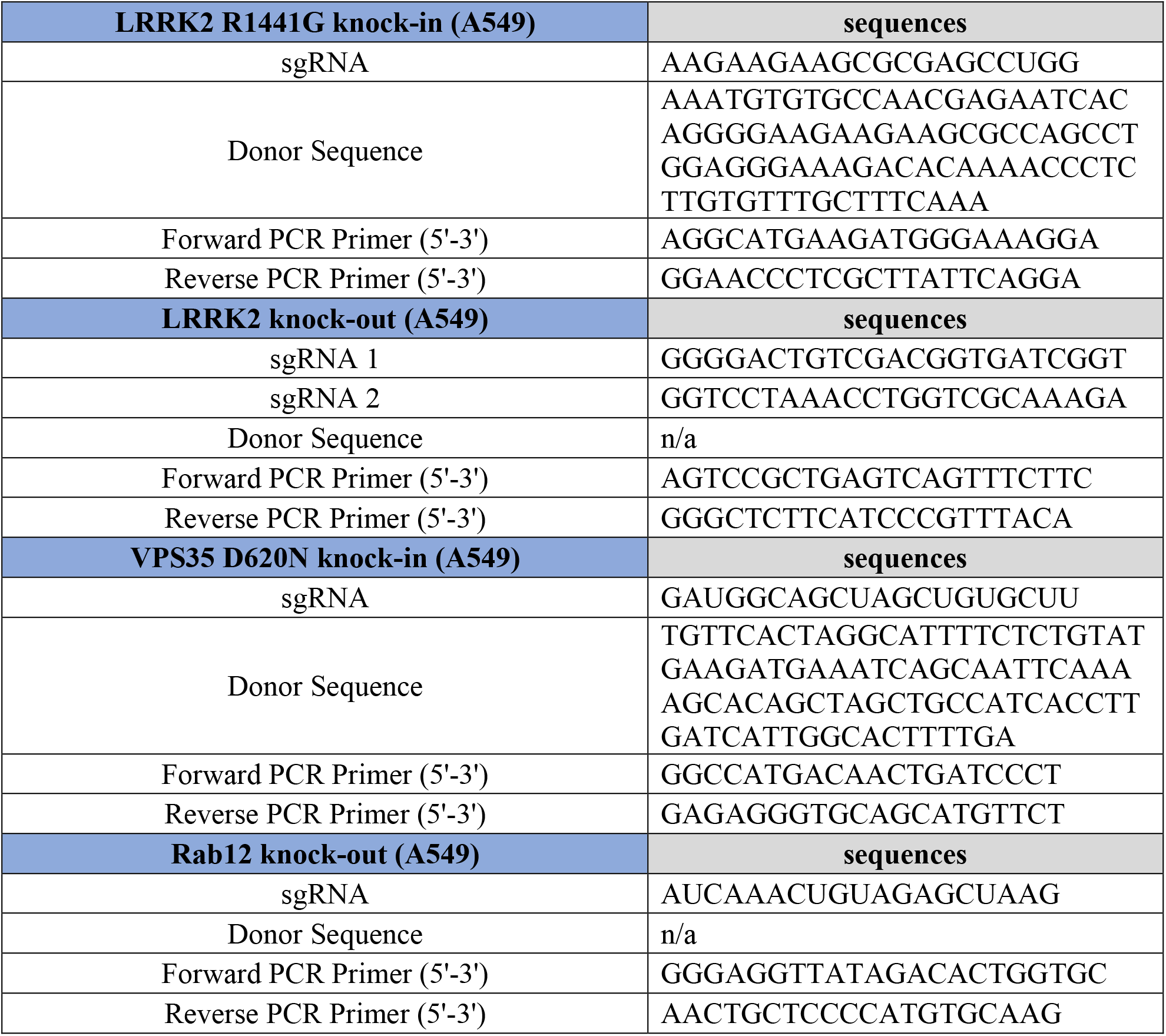

To enable the rapid isolation of lysosomes using immunopurification, A549 cells (wild type, LRRK KO, LRRK2 R1441G, RAB12 KO and VPS35 D620N) cells were transduced with lentivirus carrying the transgene cassette for expression of TMEM192-3x-HA. Stably-expressing cells were selected using resistance to Hygromycin B (Thermo Fisher Scientific, Waltham, MA, #10687010) supplied in growth medium at 200μg/mL for 21 days. Following selection, cells were screened for the stable expression of TMEM192-3x-HA in lysosomes by quantifying the percentage of cells with co-localization of anti-HA and anti-LAMP1 by immunofluorescence, and by monitoring cell lysates for expression TMEM192-3x-HA (~30 kDa) by western blot.

### Antibodies

The following primary antibodies were used for immunocytochemistry and western blot experiments:

| Target | Host species | Vendor | Catalog | Diution | Application |
| --- | --- | --- | --- | --- | --- |
| pT73 Rab10 | Rabbit | Denali | 19-4 | 1:500 | Western blot |
| pT73 Rab10 | Rabbit | Abcam | ab241060 | 1:100 | ICC |
| Rab10 | Mouse | Abcam | ab104859 | 1:500 | Western blot |
| Rab10 | Rabbit | CST | 8127 | 1:100 | ICC |
| pS106 Rab12 | Rabbit | Abcam | ab256487 | 1:500 | Western blot |
| Rab12 | Rabbit | Proteintech | 18843-1-AP | 1:500 | Western blot |
| Rab8a | Rabbit | Abcam | ab188574 | 1:500 | Western blot |
| HA | Rabbit | CST | 3724 | 1:2000 | Western blot |
| Gal3 | Mouse | BD | 556904 | 1:500 | Western blot |
| LAMP1 | Rabbit | CST | 9091 | 1:1000 | Western blot |
| LAMP1 | Mouse | Abcam | ab25630 | 1:1000 (WB)<br>1:100 (ICC) | Western blot/ICC |
| LRRK2 | Mouse | UC Davis/NIH<br>NeuroMab<br>Facility | N241A/34 | 1:500 | Western blot |
| pS935 LRRK2 | Rabbit | Abcam | ab133450 | 1:500 | Western blot |
| GAPDH | Mouse | Abcam | ab8245 | 1:5000 | Western blot |
| Actin | Mouse | Sigma-Aldrich | A2228 | 1:5000 | Western blot |

For ICC, the following secondary antibodies (Thermo Fisher) were used at 1:1000 dilution: goat anti-mouse Alexa-Fluor 488 (A32723), goat anti-rabbit Alexa-Fluor 568 (A11036)

For WB, the following secondary antibodies (LI-COR Biosciences, Lincoln, NE) were used at 1:20,000 dilution: IRDyes 800CW donkey anti-rabbit IgG (#926-32213), 680RD donkey antimouse IgG (#926-68072).

### siRNA-mediated KD of LRRK2 and Rab GTPases

A549 cells were transfected with Dharmacon SMARTpool siRNA targeting 14 Rab GTPases, LRRK2 and non-targeting scramble control (Horizon Discovery, Cambridge, United Kingdom), using DharmaFECT 1 (Horizon, T-2001-01). Cells were collected 3 days after transfection for protein or mRNA analysis.

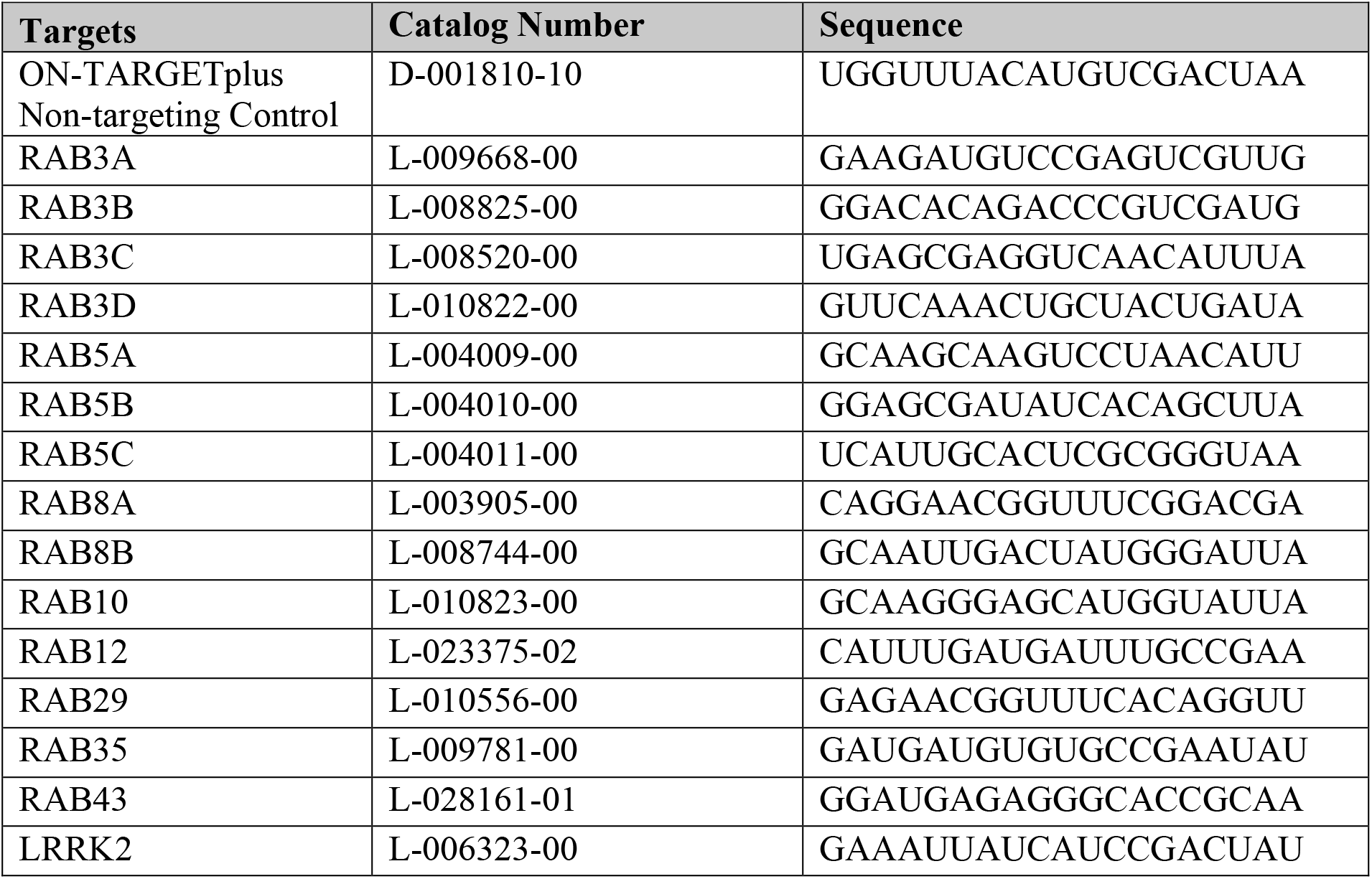

### RT-PCR-based analysis of Rab expression

The total RNA was extracted from cells using RNeasy Plus Micro Kit (QIAGEN, Hilden, Germany, #74034). cDNA was synthesized from 1~2 μg of RNA using Superscript IV VILO master mix (Thermo Fisher #11756050). The cDNA was diluted 3-fold and 1 μl of diluted cDNA was used as template. To measure the relative expression levels of mRNAs by RT-qPCR, Taqman Fast Advanced Master Mix (Thermo Fisher #4444557) was used, together with genespecific primers using TaqMan Assays (Thermo Fisher). GAPDH was used as the housekeeping gene. The PCR reaction was run using QuantStudio™ 6 Flex Real-Time PCR System, 384-well (Thermo Fisher). Gene expression was analyzed using 2^(delta-delta Ct) method with GAPDH as internal controls.

| Taqman assay ID | Gene name | Dye |
| --- | --- | --- |
| Hs00923221_m1 | RAB3A | FAM-MGB |
| Hs01001137_m1 | RAB3B | FAM-MGB |
| Hs00384846_m1 | RAB3C | FAM-MGB |
| Hs00758197_m1 | RAB3D | FAM-MGB |
| Hs00702360_s1 | RAB5A | FAM-MGB |
| Hs05027271_g1 | RAB5B | FAM-MGB |
| Hs00904926_g1 | RAB5C | FAM-MGB |
| Hs00180479_m1 | RAB8A | FAM-MGB |
| Hs00213006_m1 | RAB8B | FAM-MGB |
| Hs00794658_m1 | RAB10 | FAM-MGB |
| Hs01391604_m1 | RAB12 | FAM-MGB |
| Hs01026316_m1 | RAB29 | FAM-MGB |
| Hs00199284_m1 | RAB35 | FAM-MGB |
| Hs03006628_gH | RAB43 | FAM-MGB |
| Hs01115057_m1 | LRRK2 | FAM-MGB |
| Hs99999905_m1 | GAPDH | VIC |

### MSD-based analysis of pT73 Rab10, total and pSer935 LRRK2

LRRK2, pS935 LRRK2 and pT73-Rab10 MSD assays were previously established (23). Briefly, capture antibodies were biotinylated using EZ-Link™ NHS-LC-LC-Biotin (Thermo Fisher, #21343), and detection antibodies were conjugated using Sulfo-TAG NHS-Ester (Meso Scale Diagnostics (MSD), Rockville, MD, R31AA-1). 96-well MSD GOLD Small Spot Streptavidin plates (MSD, L45SSA-1) were coated with 25 μl of capture antibody diluted in Diluent 100 (MSD, R50AA-2) for 1 h at room temperature with 700 rpm shaking. After three washes with TBST, 25 μl samples were added each well and incubated at 4 °C overnight with agitation at 700 rpm. After three additional washes with TBST, 25 μl of detection antibodies were added to each well diluted in TBST containing 25% MSD blocker A (MSD, R93AA-1) together with rabbit (Rockland Immunochemicals, Pottstown, PA, D610-1000) and mouse gamma globin fraction (Rockland, D609-0100). After a one hour incubation at room temperature at 700 rpm and three washes with TBST, 150 μl MSD read buffer (MSD R92TC, 1:1 diluted with water) was added, and plates were read on the MSD Sector S 600.

| Assay | Antibody type | targets | Vendor | Cat No. | Concentration (µg/mL) |
| --- | --- | --- | --- | --- | --- |
| pS935 LRRK2 | Capture | pS935 LRRK2 | Abcam | ab133450 | 0.5 |
|  | Detection | Total LRRK2 | BioLegend | 808201 | 1 |
| Total LRRK2 | Capture | Total LRRK2 | BioLegend | 844401 | 0.5 |
|  | Detection | Total LRRK2 | BioLegend | 808201 | 1 |
| pT73 Rab10 | Capture | pT73 Rab10 | Denali | 19-4 | 1 |
|  | Detection | Total Rab10 | Abcam | ab181367 | 2 |

### Cell lysis and immunoblotting

Cells were lysed in lysis buffer (Cell Signaling Technology (CST), Danvers, Massachusetts, USA, #9803) supplemented with cOmplete tablet (Roche, Penzburg, Germany, #04693159001), phosSTOP (Roche #04906837001) and Benzonase nuclease (Sigma-Aldrich, St. Louis, MO, E1014). Cell lysates were prepared by incubating with NuPage LDS Sample Buffer (Thermo Fisher, NP0007) and NuPAGE™ Sample Reducing Agent (Thermo Fisher, NP0004) for 5 min at 95 °C to denature samples. Lysates were loaded onto NuPAGE 4-12% Bis-Tris gels (Thermo Fisher). Proteins were transferred to nitrocellulose membranes (Bio-Rad, Hercules, CA) using Trans-Blot Turbo Transfer System (Bio-Rad). Membranes were blocked with Rockland blocking buffer at room temperature for one hour (Rockland Immunochemicals, Pottstown, PA), incubated with primary antibody (diluted in Blocking Buffer) overnight at 4 °C, and then with secondary antibodies (1:20,000 diluted in Blocking Buffer, LI-COR) for one hour at room temperature. Odyssey CLx Infrared Imaging System (LI-COR) was used for western blot detection and quantitation.

### Cell culture and treatment

HEK293 cells and A549 cells were cultured in DMEM media (Thermo Fisher #11965-092) containing 1% Pen/Strep and 10% FBS (VWR International, Radnor, PA, #97068-085). For LLOMe treatment, LLOMe (Sigma-Aldrich, #L7393) was added at 1 mM for 2 hours or 4 hours prior to fixation or lysing cells for downstream analysis.

### Generation of Dox-inducible cell lines expressing Rab12

Doxycycline-inducible cell lines were generated to stably express WT Rab12 or a phosphodeficient mutant of Rab12 (S106A) in RAB12 KO A549 cells. Briefly, lentiviral constructs were generated by cloning 3X FLAG-RAB12 (or RAB12 S106A) into the pLVX-TetOne-Puro vector. Lentivirus was produced by transfecting the plasmids in HEK293T cells using Lenti-X™ Packaging Single Shots (Takara Bio, Kusatsu, Shiga, Japan, #631278). The media containing lentivirus were collected from transfected cells and were further concentrated by 50-fold using Lenti-X Concentrator (Takara, #631231). RAB12 KO A549 cells were infected with lentivirus expressing WT 3XFLAG-RAB12 or 3XFLAG-RAB12 S106A mutant. Cells carrying the lentiviral vectors were selected with puromycin (1 μg/ml). To enable the expression of WT Rab12 or the Rab12 S106A mutant, doxycycline (0.1, 0.5 and 1 μg/ml) was added in the cell culture for 3 days.

### Immunoprecipitation of lysosomes using TMEM192-HA ^x3^

Lysosomes were isolated by immunoprecipitation from cells expressing the TMEM192-HA^x3^ transgene as described previously(28) with the following modifications. Cells were plated in 15-cm culture dishes such that they reached full confluency on the day of the experiment. All subsequent steps were performed at 4°C or on ice with pre-chilled reagents, unless otherwise noted. Media was removed and monolayers were rinsed with KPBS buffer (136mM KCl, 10mM KH_2_PO_4_, pH 7.25), harvested by scraping into fresh KPBS and pelleted via centrifugation. Cell pellets were resuspended in KPBS+ buffer (KPBS supplemented with 3.6% (w/v) iodixanol (OptiPrep; Sigma-Aldrich), cOmplete protease inhibitor (Roche), and PhosStop phosphatase inhibitor (Roche)), and cells were fractionated by passing the suspension through a 21g needle 5 times followed by centrifugation at 800g for 10min. Post-nuclear supernatant (PNS) was harvested and incubated with anti-HA magnetic beads (pre-blocked with BSA and washed with KPBS buffer; Thermo Fisher) for 15min with end-over-end rotation. Lysosome-bound beads were washed 3 times with KPBS+ buffer, and samples used for immunoblotting were eluted from beads by heating to 95°C for 10min in 1x NuPAGE LDS Sample Buffer (Thermo Fisher).

### Analysis of pRab10 levels on isolated lysosomes from WT, RAB12 KO, and PD-linked variant KI A549 cell models

For analysis of pRab10 levels on isolated lysosomes, one confluent 15cm plate of cells was used per experimental condition. Cells were treated with 1 mM LLOMe (or vehicle) for 2h at 37°C prior to isolation of lysosomes via anti-HA immunoprecipitation as described above. Lysosome immunoprecipitations were performed with 60 μl of anti-HA magnetic bead slurry per condition. Immunoblotting for pRab10 and pRab12 levels was performed in parallel with analysis of total Rab10 and Rab12 levels, as detailed above, using 20% of total immunoprecipitated material per condition. pT73 Rab10, pS106 Rab12, total Rab10, total Rab12, and HA band intensities were quantified from immunoblots using ImageStudio Lite software (LI-COR), and the phospho- and total Rab band intensities were normalized to HA band intensity within each experimental condition. Data were normalized to the median value within each replicate and was then normalized to the mean value of vehicle-treated WT samples across replicates.

### Analysis of LRRK2 levels on isolated lysosomes from A549 cell models

For analysis of LRRK2 levels on isolated lysosomes, three confluent 15cm plates of cells (seeded 24h prior to the assay start) were used per experimental condition. Cells were treated with 1 mM LLOMe (or vehicle) for 4h at 37°C and then lysosomes were isolated via anti-HA immunoprecipitation as described above. Lysosome immunoprecipitations were performed with 150 μl of anti-HA magnetic bead slurry per condition. For immunoblot detection of endogenous LRRK2, 25% of the total immunoprecipitated material (per condition) was loaded onto a 3-8% Tris-Acetate gel (Thermo Fisher), fully resolved gels were transferred to nitrocellulose membranes, probed overnight at 4°C with a 1/500 dilution of mouse anti-LRRK2 (clone N241A/34; UC Davis/NIH NeuroMab Facility, Davis, CA), and imaged using standard immunoblotting protocol detailed above. LRRK2 and HA band intensity was quantified from immunoblots using ImageStudio Lite software (LI-COR), LRRK2 intensity was normalized to HA band intensity within each experimental condition, data was normalized to the median value within each replicate, and then was normalized to the mean value of vehicle-treated WT samples across replicates.

### Immunostaining of pT73 Rab10 and Rab10, and imaging analysis

WT, RAB12 KO and LRRK2 KO A549 cells were seeded in 96 well plates (PerkinElmer, Waltham, MA, Phenoplate, #6055302), and then treated with vehicle or LLOMe (1 mM). After 2 hours, cells were fixed with 4% PFA for 15 mins, permeabilized and blocked with blocking buffer (5% Normal Donkey Serum/ 0.05% Triton/PBS) for 1 hour at room temperature. Primary antibodies were diluted in blocking buffer and incubated overnight at 4 °C. After 3 washes with PBS/0.05% Triton, secondary fluorescently-labeled antibodies were diluted in blocking buffer and incubated for 1 hour at room temperature. DAPI (1:1000) and cell mask deep red (1:5000, Thermo Fisher, C10046) were diluted in PBS/0.05% Triton and incubated for 10 min. After 3 washes with PBS/0.05% Triton, the cell plates were imaged on an automated confocal high-content imaging system (PerkinElmer, Opera Phenix Plus High-Content Screening System) using a 63X water immersion objective lens with excitation lasers (405 nm, 488 nm; 561 nm; 640 nm) and preset emission filters. Channels were separated to avoid fluorescence crosstalk. A custom analysis was developed in the Harmony 4.9 image analysis software (PerkinElmer) to enable image analysis. For analysis of puncta intensity, pT73 Rab10 or total Rab10 spots were defined using “Finding Spots” building blocks, and the sum of corrected spot intensity per cell was used to measure puncta signals. The colocalization between pT73 Rab10 puncta and LAMP1-positive lysosomes were measured with object-based analysis. Briefly, pT73 Rab10 and LAMP1 spots were independently defined using separate “Find Spots” building blocks. Colocalized pT73 Rab10 and LAMP1 spots were identified using the geometric center overlap method within the “Select Population” tool. The average number of colocalized spots were calculated per cell from 16 fields per well and averaged across the well.

### Live cell imaging of Rab12 and LRRK2 in HEK293 cells

HEK293 cells were transfected with eGFP-LRRK2 and mCherry-Rab12 plasmids (pcDNA3.1 vectors) using Lipofectamine LTX with Plus Reagent (Thermo Fisher #15338100), and cells were plated onto poly-lysine coated 96-well plates (Corning Inc. Corning, NY, BioCoat plates, #354640). 2 days after transfection, cells were incubated with Hoechst 33342 (1 μg/ml, Thermo Fisher #62249) and CellMask™ Deep Red Plasma membrane Stain (1:2000, Thermo Fisher C10046) for 10 min. After replacing the cell culture media containing 1 mM LLOMe, the cell plates were immediately started for live cell imaging on an automated spinning-disk confocal high-content imaging system (Perkin Elmer, Opera Phenix) using a 40X water immersion objective lens under 5% CO_2_ and 37 °C condition. The confocal images were taken every 10 min for 90 min in total.

### Time lapse cell segmentation analysis

Fields of view containing cells co-transfected for eGFP-LRRK2 and mCherry-Rab12 were manually selected from the time lapse dataset across three independent experiments. For each field, channel, and timepoint, the z-stack was converted to a 2D image by maximum intensity projection, then the background intensity was estimated by smoothing the image with a Gaussian filter with a kernel standard deviation of 50 pixels (~14.8 μm) using the ndimage module in scipy v1.9.3(33). The background was subtracted from the original image and all pixels below the background intensity were set to zero. Background subtracted images were loaded into napari v0.4.17(34) as 2D+time images for segmentation.

To better visualize cells co-expressing low levels of LRRK2 and Rab12, the contrast limits were set to between 0 and 125 AU for LRRK2 and between 0 and 600 AU for Rab12. Cells were included in the segmentation if they 1) appeared morphologically healthy, 2) were visible throughout the time series, and 3) co-expressed LRRK2 and Rab12 at levels above background, but below the maximum contrast limit. Cells were segmented using the brush tool in napari using a brush size of 10 pixels (~3.0 μm). Cells were painted to the edge of the cell border including the nucleus but excluding signal from cell debris and other extracellular sources. Tracking cells across frames was typically possible through a combination of proximity and morphology, but where the assignment was ambiguous, those cells were excluded from further analysis (n=2). The resulting dataset contained 55 segmented cells.

For each cell, the non-background subtracted LRRK2 and Rab12 signals were extracted and normalized to between 0.0 and 1.0 by subtracting 200 AU and then dividing by 800 AU. Values above 1.0 or below 0.0 were set to 1.0 or 0.0 respectively. For all pixels under the cell mask, the Pearson correlation coefficient I was calculated between the normalized LRRK2 and Rab12 signals using the pearsonr function in the scipy.stats module. Cell properties such as area, perimeter, and mean intensity in each channel were extracted for each timepoint using the regionprops function in scikit-image v0.19.3(35). Cells were filtered for quality by fitting a least squares line to the mean intensity of both LRRK2 and Rab12 signal for each and excluding any cells where the slopes were negative (n=31 cells excluded), resulting in 24 validated cell traces. Normalized intensity and correlation coefficients were plotted as mean of all 24 traces +/- standard error (SEM) using Prism v9.3.0 (GraphPad).

### Statistical Analysis

Data are shown as means ± SEM, and all statistical analysis was performed in GraphPad Prism 9. Unpaired (or paired) t-tests were used for statistical analyses of experiments with two treatment groups. For more than two groups, analysis was performed using one-way analysis of variance (ANOVA) with Tukey’s multiple comparison, one-way ANOVA with Sidak’s multiple comparison test, one-way ANOVA with Dunnett’s multiple comparison test, repeated measures (RM) one-way ANOVA with Dunnett’s multiple comparison or two-way ANOVA with Sidak’s test, as indicated in figure legends. Comparisons were considered significant where p <0.05. *, p< 0.05; **, p< 0.01; ***, p< 0.001; ****, p< 0.0001.

## DATA AVAILABILITY

The data that support the findings of this study are available from the corresponding author upon reasonable request.

## ACKNOWLEDGEMENTS

We thank Dara Leto and Meredith Calvert for support and guidance for imaging-based analysis and members of the Denali postdoc program and lysosomal function pathway team for useful discussions and feedback.

